# Experimental evidence of memory-based foraging decisions in a large wild mammal

**DOI:** 10.1101/2020.05.23.112912

**Authors:** Nathan Ranc, Paul R. Moorcroft, Federico Ossi, Francesca Cagnacci

## Abstract

Many animals restrict their movements to a characteristic home range. This pattern of constrained space-use is thought to result from the foraging benefits of memorizing the locations and quality of heterogeneously distributed resources. However, due to the confounding effects of sensory perception, the role of memory in home range movement behavior lacks unequivocal evidence in the wild. Here, we analyze the foraging decisions of a large mammal during a field resource manipulation experiment designed to disentangle the effects of memory and perception. Using a cognitive movement model, we demonstrate that roe deer (*Capreolus capreolus*) rely on memory, not perception, to track the spatio-temporal dynamics of resources within their home range. Our findings show a memory-based spatial transition model parametrized with experimental data can successfully be used to quantify cognitive processes and to predict how animals respond to resource heterogeneity in space and time.

## Introduction

Many animals, both territorial and non-territorial, constrain their movements to a characteristic home range, an area that is typically much smaller than their movement abilities would allow (Burt 1943). The ubiquity of this space-use pattern suggests that home ranges are adaptive and that a general mechanism underpins their emergence (Börger *et al.* 2008). In particular, home ranges are thought to result from the foraging benefits provided by *spatial memory* (Spencer 2012) – the process by which animals encode spatial relations (Fagan *et al.* 2013).

The role of spatial memory is particularly relevant when resources are spatially heterogeneous and temporally dynamic (Mueller & Fagan 2008), making foraging a complex, spatio-temporal problem. Classic ecological theory such as optimal foraging (Charnov 1976) and the ideal free distribution hypothesis (Fretwell & Lucas 1969), as well as resource selection analyses (Boyce & McDonald 1999), assume that animals have either no knowledge (a random forager) or perfect knowledge of the spatio-temporal patterns of resources (an omniscient forager). However, animal foraging decisions must rely, in fact, on imperfect information (McNamara & Houston 1985, 1987) gained through two separate streams: sensory perception, and memories of previous experiences at locations beyond the individual’s current perceptual range (Shettleworth 2001; Fagan *et al.* 2013).

In this context, memory should be adaptive whenever retaining past, site-specific information to predict the future occurrence and quality of key resources is more efficient than foraging relying on proximal mechanisms such as area-restricted search and perception (Spencer 2012). Accordingly, foragers may not only benefit from memorizing spatial locations, but also from tracking the profitability of previously visited resources by means of an *attribute memory* (Fagan *et al.* 2013). Such dynamic learning allows the forager to develop an expectation of resource quality from previous experience (Garber & Paciulli 1997) and implies the discounting of old information (Bush & Mosteller 1951; Killeen 1981). In support of this argument, theoretical studies have demonstrated the foraging advantage of memory in spatially-heterogeneous, predictable landscapes (Van Moorter *et al.* 2009; Bracis *et al.* 2015; Riotte-Lambert *et al.* 2015).

Empirically, the benefits of memory for resource acquisition have been documented for several kinds of central-place foragers, in particular hummingbirds (González-Gómez *et al.* 2011), food caching birds (Clayton & Dickinson 1998) and bumblebees (Burns & Thomson 2006). Experimental evidence of memory-based foraging decisions in wild mammalian home ranges has been instead limited (Janson 1998). The influence of memory and perception on the movement behavior of mammals has been inferred in several observational studies (Normand *et al.* 2009; Merkle *et al.* 2014, 2019; Avgar *et al.* 2015; Bracis & Mueller 2017); however, quantifying their respective influences on foraging decisions is challenging because both memory and perception can give rise to long-distance, goal-oriented movements. Field experiments have the potential to address this limitation by providing the level of control required to disentangle the effects of memory and perception (Garber & Paciulli 1997; Fagan *et al.* 2013; Bracis & Mueller 2017). In a rare field experiment in mammals, Janson showed that the home range movements of a brown capuchin (*Cebus apella*) troop deviated from a perception-based movement model (Janson 1998, 2016); however, a mechanistic, quantitative understanding of how memory affects mammalian foraging movements is still lacking.

In this study, we address this gap by formulating a memory-based model of spatial transitions to: (1) characterize and quantify the cognitive processes involved in the foraging decisions of a large mammal; and (2) predict the observed patterns of response to a resource manipulation experiment. We performed our experiment on European roe deer (*Capreolus capreolus*), a particularly well-suited species for studying the interplay between cognitive processes and resource dynamics. As browsers with limited fat reserves (Andersen *et al.* 2000), roe deer exhibit a tight association between movement and resource dynamics (Ranc *et al.* 2019), particularly during the winter months (i.e., at the time of our experiment) when food scarcity limits foraging performance and the movements of roe deer are not affected by territoriality (Liberg *et al.* 1998). In addition, because roe deer are solitary (Hewison *et al.* 1998), their foraging decisions are expected to be primarily based on individual, personal information (Dall *et al.* 2005).

Roe deer were fitted with GPS telemetry collars at a site in the Eastern Italian Alps and followed their movements during a transitory alteration of high-nutritional food accessibility at supplemental feeding sites, located within their home range (n = 18 individuals, for a total of 25 animal-years; see *Methods*). The six-week experiment, conducted over three years, consisted of three two-week phases – pre-closure, closure and post-closure – and was designed to disentangle the effects of memory and perception. During the closure phase, the food at the most-attended feeding site of each individual (hereafter referred to as Manipulated, M) was rendered inaccessible by installing a physical barrier while maintaining food presence at the site (Fig. 1b). This ensured that sensory information on resource availability remained unaltered by the manipulation.

**Figure 1.**
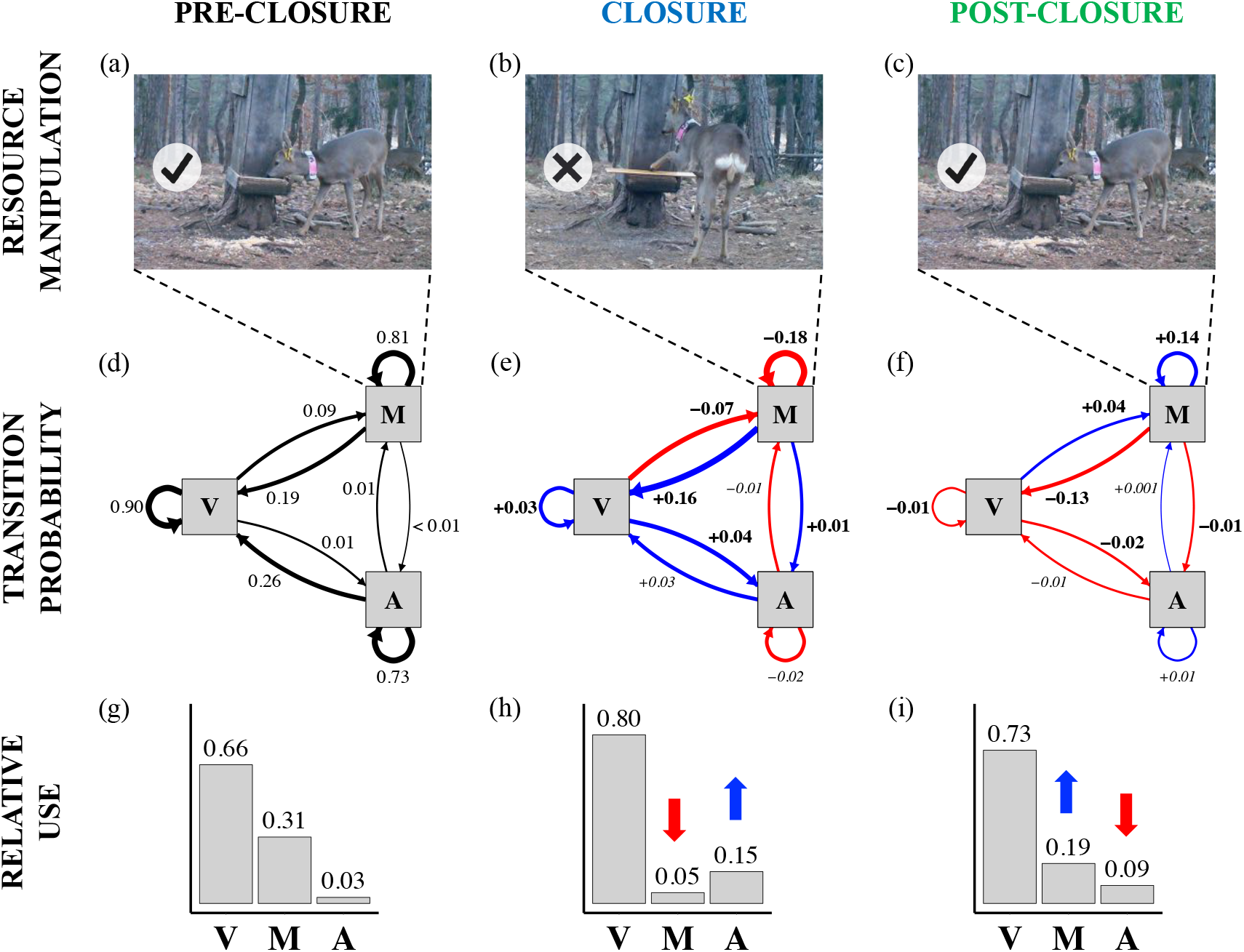
Transitory changes in resource use patterns during the experiment. Panels a-c: schematic representation of the experiment. High nutritional food is accessible at manipulated feeding sites (M) during the pre- and post-closure phases (panels a and c, respectively), and present, but inaccessible during the closure phase (panel b). Food is also present throughout the experiment at alternate feeding sites (A). Roe deer can also access natural vegetation (V). Panels d-f: transition probabilities among the three resource types – V, M and A – for pre-closure (as rates; n = 9045 transitions; panel d), closure (as net changes respect to pre-closure; n = 9187; panel e) and post-closure (as net changes respect to closure; n = 8417; panel f). For the net changes, the color of the vertices indicates a decreased or increased probability (red and blue, respectively; significant changes are in bold). Panels g-i: corresponding relative resource use, with vertical arrows illustrating the compensation pattern observed between the use of M and A.

The temporal dynamics of roe deer foraging patterns during the experiment were characterized by quantifying the fraction of time each individual spent in the vicinity of their Manipulated feeding site (M), at Alternate feeding sites available within the broader landscape (A), or in natural Vegetation (V). We then developed a model describing the transition probabilities between states (M, A, V) as a function of resource accessibility, resource preference and cognitive processes, while controlling for environmental cues (illumination patterns, temperature).

We evaluated three competing hypotheses concerning the cognitive mechanisms underlying the patterns of movement by roe deer during the experiment. (i) a null hypothesis (H1) of *omniscience-based* movement in which animals possess complete information on the spatio-temporal dynamics of resources. Accordingly, we predicted that individuals would no longer visit manipulated feeding sites when forage is inaccessible (P1.1) and respond instantaneously to actual changes in resource accessibility (P1.2), irrespective of their previous experience. (ii) a *perception-based* movement hypothesis in which animals use long-distance sensory cues to guide their foraging decisions (H2). At the spatial scale of this experiment, in which feeding sites are hundreds of meters apart, we assumed that, as in other large herbivores (Finnerty *et al.* 2017), roe deer would primarily rely on olfactory rather than visual perception because feeding sites are not visible from afar. Since the manipulation did not alter the sensory information that can be perceived at long distances we predicted that under the perception hypothesis the rate at which roe deer visited manipulated feeding sites would be constant throughout the experiment (P2.1) and that their foraging decisions should be independent of resource accessibility (P2.2). (iii) a *memory-based* movement hypothesis (H3) in which animals rely on previous experience to guide foraging decisions. We formulated a bi-component memory model consisting of a spatial memory (i.e., recollection of resource locations) and an attribute memory (i.e., recollection of the profitability at previously visited locations; *sensu* (Fagan *et al.* 2013; Merkle *et al.* 2014)). We predicted that, under this hypothesis, roe deer would decrease their visit of inaccessible feeding sites (P3.1), conditional on experienced changes in resource accessibility (P3.2). We further predicted that the influence of previously visited feeding sites on roe deer movement would slowly decrease with time since last visit (i.e., slow decay of spatial memory; P3.3) and that the expected value of feeding sites would primarily rely on very recent experience (i.e., fast decay of attribute memory; P3.4). Further details on the mathematical formulations of the three above hypotheses can be found in the *Methods* section.

## Results

### Transitory changes in resource use patterns

The experiment led to significant changes in movement rates between the three resource types (Fig. 1, panels d-f). Prior to closure, roe deer used vegetation (V), manipulated feeding site (M), and alternate feeding sites (A), at rates 66%, 31%, 3%, respectively (Fig. 1g). When in vegetation, roe deer had a 0.9 probability (per unit time) of remaining, a 0.09 probability of visiting M and a low (0.01) probability of visiting A. Closure of M (Fig. 1e) led to decreases in the probability of individuals remaining at their respective M (−0.18), and decreases in transitions from vegetation to M (−0.07) – responses that are consistent with P1.1 and P3.1 but inconsistent with P2.1. Roe deer compensated for the loss of M by increasing their movements from vegetation towards A (+0.04). Re-opening of the M sites (Fig. 1f) led to a recovery of pre-closure patterns of transition probabilities with, in particular, increases in probabilities of residence at M (+0.14) and transitions from vegetation to M (+0.04), and a decrease in transition probability from vegetation to A (−0.02).

As a result of these movement responses, resource use shifted dramatically between the different phases of the experiment (Fig. 1, panels g-i). As noted earlier, during pre-closure, roe deer primarily used V (66%), followed by M (31%) and rarely A (3%; Fig. 1g). Following closure, roe deer use of M dropped to 5% and use of A increased to 15% (Fig. 1h). Following re-opening, roe deer use of M recovered to 19% and use of A declined to 9% (Fig. 1i). While less marked, the temporal changes in use of V mirrored those of A, increasing from 66% (Fig. 1g) to 80% during closure (Fig. 1h), and then declining to 73% during post-closure (Fig. 1i).

#### Evidence for memory-based foraging decisions

The memory-based model had overwhelming stronger support (AIC = 20262; n = 26,649 movement transitions; H3) compared to the omniscience- (ΔAIC = +829; H1) and perception-based movement models (ΔAIC = +1093; H2). Overall, the estimates and confidence intervals of the parameters shared among the three models (within-state resource accessibility, preference for M, minimum daily temperature and illumination) were highly consistent (Fig. 2), suggesting that the differences in model support result from the differences in the underlying cognitive formulations (rather than spurious correlation with other covariates affecting the probability of movement).

**Figure 2.**
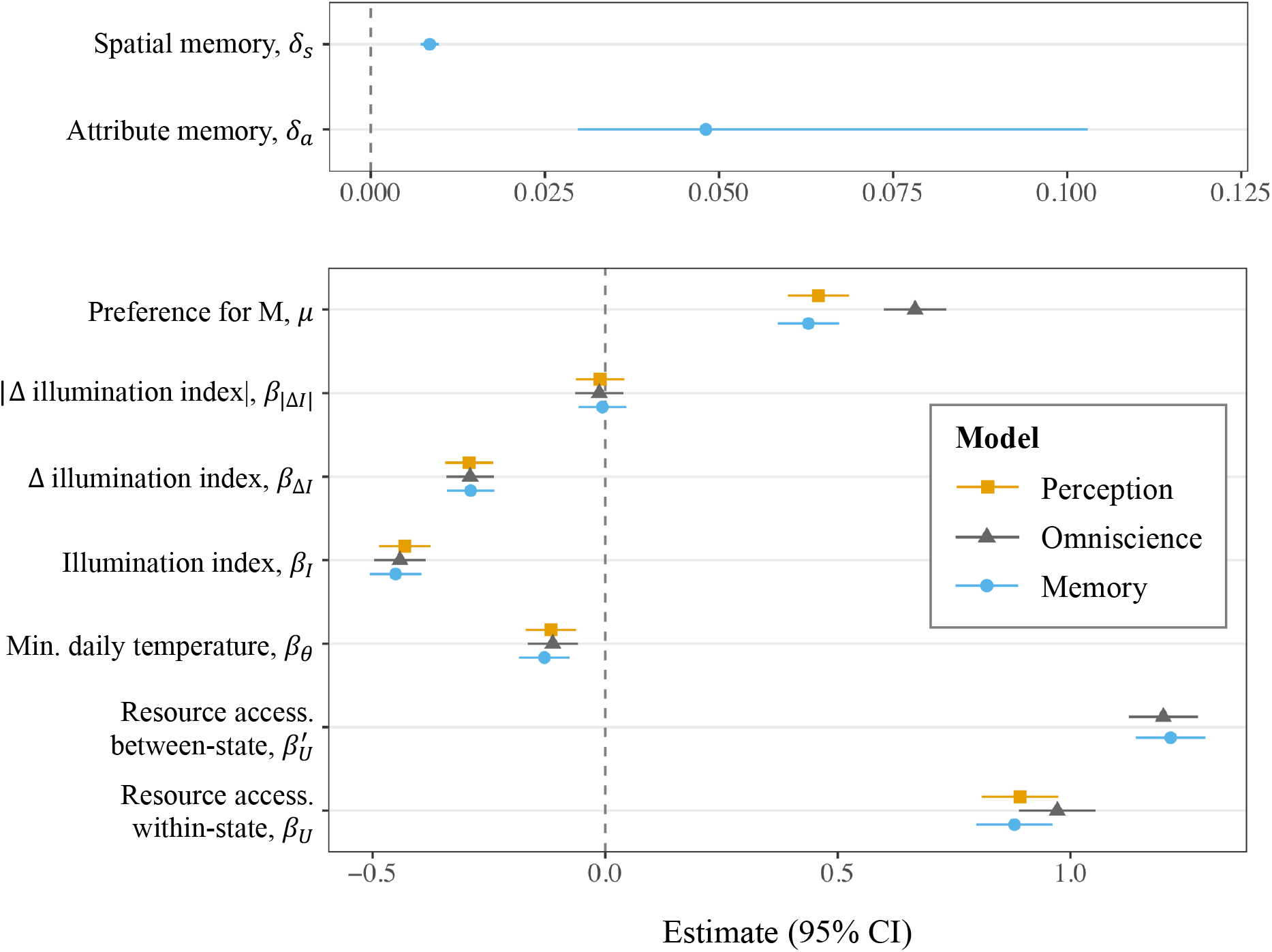
Parameter estimates. The estimates for the perception-based (orange square), omniscience-based (grey triangle) and memory-based (blue circle) models are plotted with the corresponding 95% marginal confidence intervals. Memory parameters are presented separately for readability (different magnitudes).

Figure 3 illustrates the predictive capabilities of the memory-, omniscience- and perception-based movement models by showing the temporal trends in the probability of moving from vegetation to either the manipulated feeding site (i.e., V-to-M transition, black lines) or alternate feeding sites (i.e., V-to-A transition; red lines). Specifically, the memory-based model captures both the sudden drop in the visiting probability of the manipulated FS following experimental closure (Fig. 3a, black line) and the respective compensatory increase in the visitation of alternate FS (Fig. 3a, red line). The omniscience-based model also predicts a decline in V-to-M transitions following closure (Fig. 3b, black line); however, it fails to capture the compensatory increase in V-to-A transitions following closure, and the respective upward and downward temporal trends in the probabilities of V-to-A and V-to-M transitions during the closure phase that are captured by the memory-based model (compare red and black lines in Fig. 3, panels a and b, during the closure period). Similarly, the memory-based model is the only model that captures the downward and upward temporal trends in the probabilities of V-to-A and V-to-M transitions during the post-closure period (compare red and black lines in Fig. 3, panels a and b, during the post-closure period). Finally, the perception-based model fails to capture any of the temporal shifts in foraging behavior that occur during the experiment (Fig. 3c).

**Figure 3.**
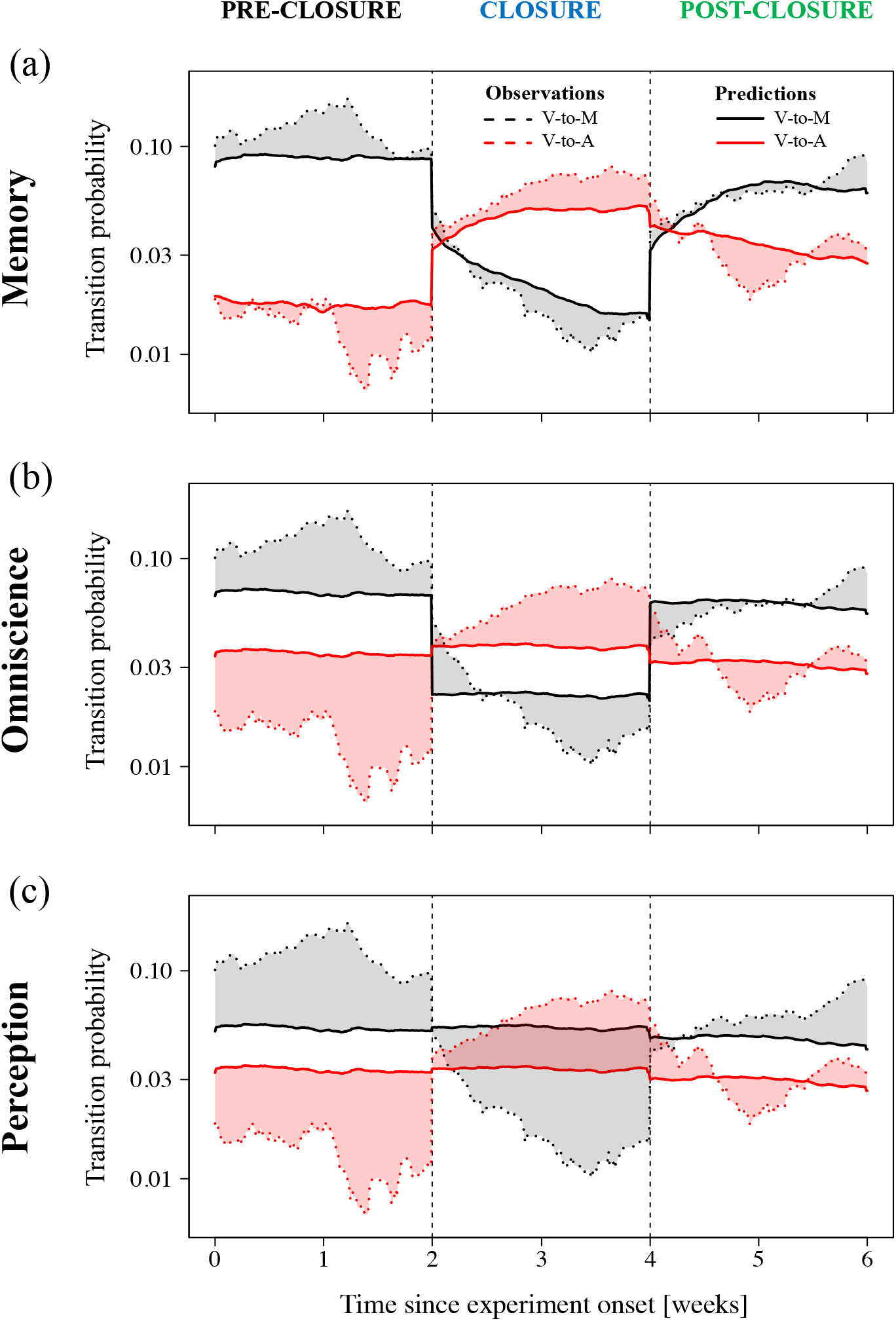
Predictions from the three competing cognitive models. The predicted (solid lines) and observed (dotted lines) transition probabilities from vegetation (V) to either manipulated feeding site (M; V-to-M transition; black lines) or alternate feeding sites (A; V-to-A; red lines) during the three experimental phases are plotted as running four-day means across all animal-years on a log-scale. Transition probabilities were calculated from 5941 transitions (from V to either V, M or A) during pre-closure, 7336 during closure and 6107 during post-closure. The red and grey shading in each panel indicate the difference between the predicted and observed probabilities of V-to-M and V-to-A transitions respectively.

Spatial memory was the most important variable influencing roe deer selection of distant resources (δ_s_; ΔAIC = +500; Table 1), and the main driver underlying the higher support of the memory-over omniscience-based movement models. Roe deer favored recently-visited resources (P3.3): spatial memory decreased exponentially with time since last visit with a half-life (t_1/2_) of 3.4 days (δ_s_=8.5 ×10^−3^ h^−1^; 95% CI: 7.2-9.8 ×10^−3^; Fig. 4).

**Table 1.**
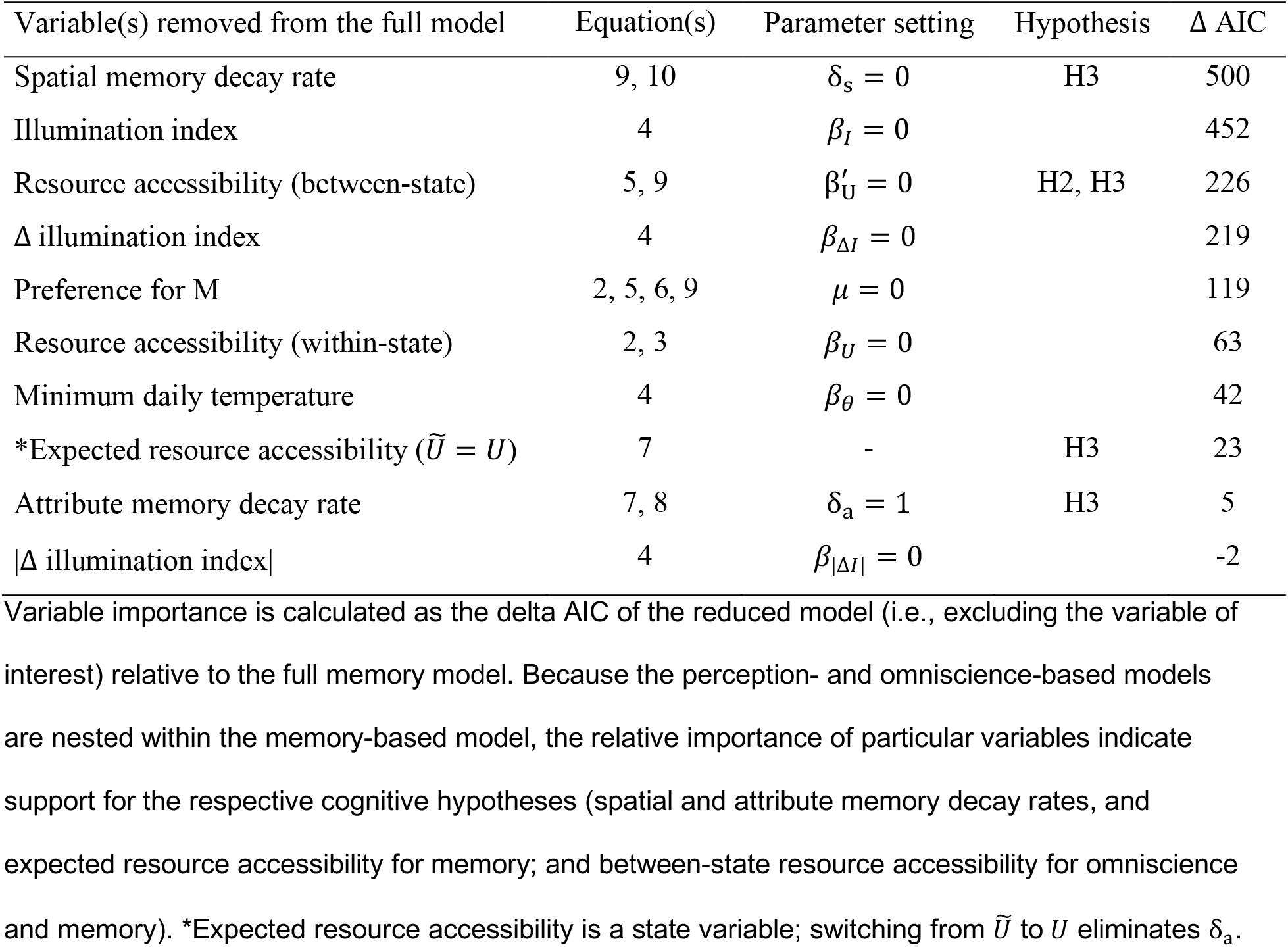
Variable contributions to the memory-based model.

**Figure 4.**
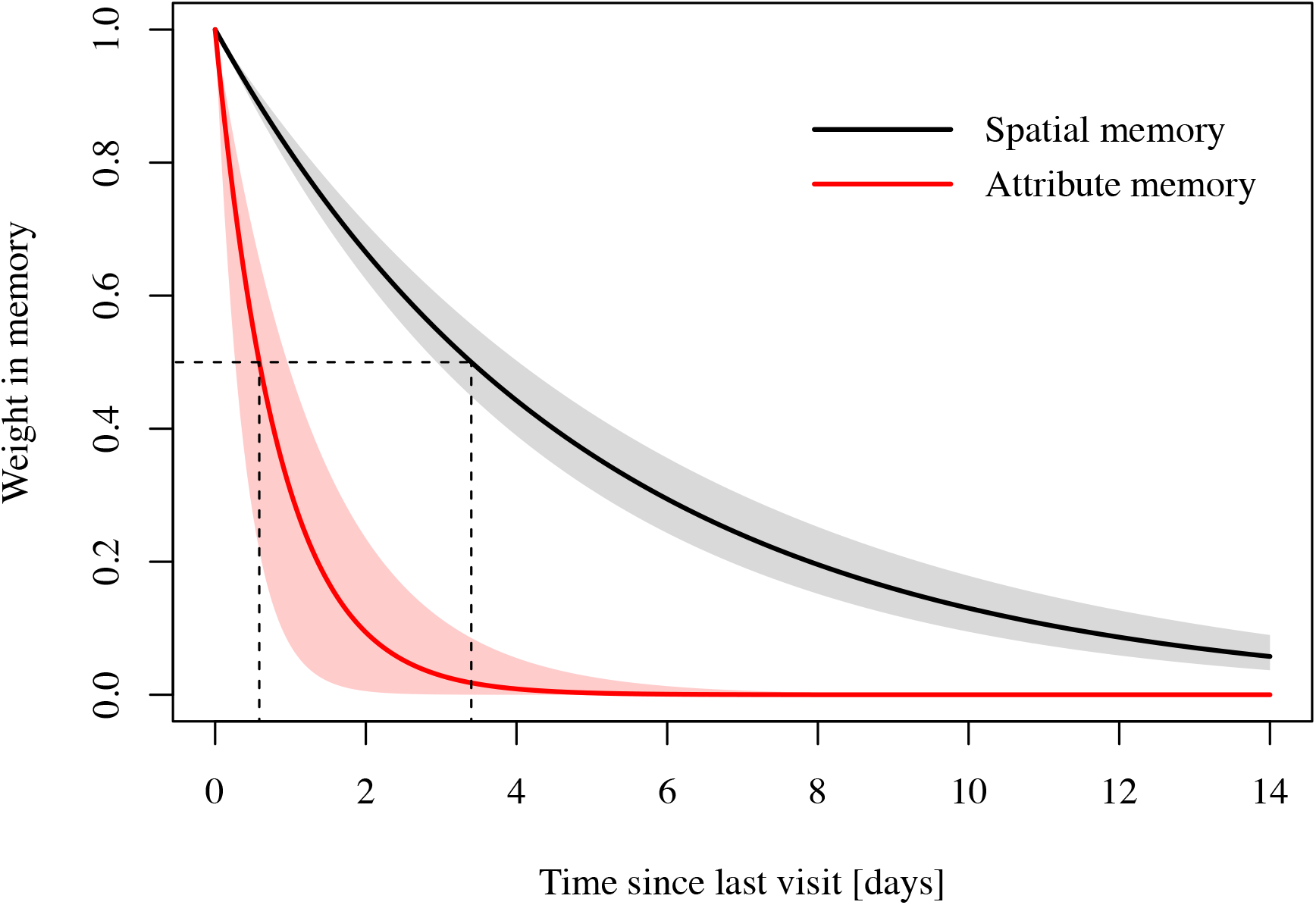
Decay of spatial and attribute memory with time since last visit. Shaded areas indicate 95% marginal confidence intervals, and dashed lines represent half-life values (t_1/2_). Spatial memory decreased exponentially with time since last visit at rate 8.5 ×10^−3^ h^−1^ (95% CI: 7.2-9.8 ×10^−3^; t_1/2_ = 3.4 days), and attribute memory decays at rate 4.8×10^−2^ h^−1^ (CI: 3.0-10.3 ×10^−2^; t_1/2_ = 0.58 days).

When evaluating the profitability of distant resources (i.e., between-state), roe deer strongly selected for accessible feeding sites (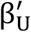; ΔAIC = +226; third variable importance; Table 1), consistent with either omniscience or memory, but contradicting the perception hypothesis (P2.2 not supported). In addition, roe deer foraging decisions were consistent with a selection for expected, rather than actual, resource accessibility (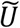; ΔAIC = +23; Table 1), thereby supporting the memory hypothesis over the omniscience hypothesis (i.e., P3.2 supported; P1.2 not supported). The rapid rate of attribute memory decay – half-life of 0.58 days (δ_a_=4.8×10^−2^ h^−1^; 95% CI: 3.0-10.3 ×10^−2^; Fig. 4) – indicates that roe deer expectations of resource profitability primarily relied on recent experience of FS accessibility (P3.4) i.e., time averaging over short periods. Support for time averaging of previous experiences, as opposed to reliance on the last experience, was relatively weak (ΔAIC = +5 when attribute memory decayed instantly i.e., δ_a_ = 1; Table 1).

Roe deer residence time at FS (as indicated by the probability of remaining at given site per unit time i.e., within-state) was also influenced by resource accessibility, with deer attending FS for significantly shorter durations when food was inaccessible (*β*_*U*_; ΔAIC = +63; Table 1). When resources were equally accessible, roe deer preferred the manipulated FS over alternate FS (*μ*; ΔAIC = +119; Table 1), leading to a higher probability of transitions from vegetation to M compared to transitions to A (Supplementary S1: Fig. S1), and to a higher residence time at M (not shown).

Environmental conditions also influenced roe deer foraging behavior during the experiment (Table 1). In particular, roe deer use of FS was markedly affected by illumination with peak visitation rates at dusk and a higher probability of visitation at night than during the day (Supplement S1, Fig. S1), as indicated by the significant effects of illumination index and its rate of change (*β*_I_ and *β*_Δ*I*_; respectively the second and fourth most important explanatory variables in the final movement model; ΔAIC = +452 and ΔAIC = +219; Table 1). Instead, the absolute rate of change of the illumination index had a negligible effect (*β*_|Δ*I*|_; ΔAIC = −2; Table 1), and therefore was not retained in the final memory model. There was also an effect of minimum daily temperature: with roe deer attended FS more intensely on colder days (*β*_*θ*_; ΔAIC = +42; Table 1).

## Discussion

Developing a unified theory of animal space-use requires a mechanistic understanding of the cognitive processes underlying animal movement decisions, and their fitness consequences in nature (Mitchell & Powell 2004; Börger *et al.* 2008; Spencer 2012). In this study, we disentangled the respective influences of perception and memory on the foraging behavior of a large mammal by assessing the abilities of perception-, memory- and omniscience-based movement models to capture individual roe deer responses with data from an *in situ* resource manipulation experiment. As seen in Figure 3, combining a dynamic, bi-component memory model (H3) with environmental cues allowed us to accurately predict how roe deer shifted resource use in response to the experimentally-imposed shifts in resource accessibility. Instead, the mismatch between the predictions of a corresponding perception-based model (H2) and the observations indicates that the foraging decisions of roe deer during the experiment were not caused by long-distance sensory cues of resource presence. The ability of roe deer to perceive the smell of supplemental food from afar is largely unknown. It could be that roe deer are not able to perceive food presence through olfaction from afar (i.e., memory is their only source of information). Alternatively, the information encoded in their memory about a resource’s accessibility over-rode sensory cues. Such hierarchical processing of information has been shown for wild capuchin monkeys (*Cebus capucinus*), which primarily used memory, overrunning conflicting perceptual cues, when resource locations were predictable (Garber & Paciulli 1997).

In addition, by accounting for the temporal lags in the movement behavior of individuals around experimental manipulations, the predictions of the memory-based model (H3) provide a much better fit to the observations than our omniscience-based model (null; H1) that assumes perfect knowledge of the changing resource dynamics by roe deer (Fig. 3). This key result highlights the fact that animal foraging decisions are indeed based on incomplete information on the location and quality of available resources (McNamara & Houston 1985, 1987), a factor that is typically not accounted for in analyses of animal movement in the wild.

In previous observational studies, memory effects have been inferred either from the re-visitation of geographic locations (Dalziel *et al.* 2008; Wolf *et al.* 2009; Oliveira-Santos *et al.* 2016), or from the discrepancy between observations and either random or perception-based movement models (Janson 1998, 2016; Normand *et al.* 2009; Hopkins 2016). Our findings build upon these results in two important ways. First, in contrast to studies of geographic re-visitation (Dalziel *et al.* 2008; Wolf *et al.* 2009; Oliveira-Santos *et al.* 2016), the findings of our experimental study rule-out the possibility that the observed movement patterns are caused by perception rather than memory – two mechanisms that are often confounded in observational studies of animal movement (Garber & Paciulli 1997). Moreover, our results show that the impacts of memory on movement behavior are dynamic, and conditional on resulting performance – in this case, the effects of memory are mediated by the impacts on the resulting foraging success of individuals. Second, rather than inferring the influence of memory from discrepancies with random or perception-based movement behavior (Janson 1998, 2016; Normand *et al.* 2009; Hopkins 2016), in this study, we explicitly formulated a memory process and showed that it had higher support and predictive ability than corresponding perception-, and omniscience-based movement models.

Our experiment demonstrates that roe deer foraging decisions primarily rely on memory, consistent with previous enclosure-based experiments showing that large herbivores are capable of memorizing the location of available food resources (Gillingham & Bunnell 1989; Edwards *et al.* 1996; Laca 1998). The probability that roe deer visited particular resource patches decreased exponentially with time since last visit with a corresponding half-life for spatial memory of 3.4 days (Fig. 4). This finding contrasts with two recent studies showing that the movements of bison (*Bison bison*) (Merkle *et al.* 2014) and caribou (*Rangifer tarandus*) (Avgar *et al.* 2015) are influenced by long-term spatial information i.e., negligible or no decay of spatial memory over six months or more. The relatively rapid decay of spatial memory estimated in this study also has to be interpreted in the context of the species re-visitation rate of locations and resources within their home range. Given the average movement rate with respect to the home range size in the monitored population (63 m h^−1^, 28 ha biweek^−1^; (Ranc *et al.* 2019)), roe deer typically visit much of their home range in just a few days (as opposed to caribou and bison that range over much larger areas). As a consequence of this high background re-visitation rate, despite the relatively high estimated decay, roe deer spatial memory rarely, if ever, dropped to zero, i.e., feeding sites were never totally forgotten. The rapid decrease in the memory with elapsed time since last visit allows roe deer to rapidly shift away from less profitable resources, and hence enable them to quickly adapt to spatio-temporal changes in resource availability, as seen in this experiment.

From a roe deer’s perspective, the availability of food at a given feeding site is unknown, but can, to some degree, be predicted based upon the animal’s prior experiences. As time increases, old information about resource quality becomes increasingly unreliable over more recent experiences (Bush & Mosteller 1951; Killeen 1981), and therefore should be discounted at a rate commensurate with the temporal scale of environmental change (McNamara & Houston 1985, 1987). Our finding of a rapid decay in attribute memory (half-life of 0.58 day; Fig. 4) implies that roe deer primarily rely on their last experience to evaluate feeding site quality. This result is consistent with enclosure-based experiments in least chipmunks (*Tamias minimus*) and golden-mantled ground squirrels (*Spermophilus lateralis*), which suggest that individuals increasingly rely on recent experience when resource dynamics are slow (Devenport & Devenport 1994). In contrast, in a recent study of bison, Merkle et al. (2014) showed that individuals appear to rely on long-term memory of profitability to inform their selection of grazing meadows (slow decay of attribute memory in summer: half-life of 10.4 days and, negligible decay in winter). In our system, the movement transitions between resources occur over a few hundred meters (i.e., over relatively short distances compared to roe deer movement rates). Information about the profitability of resource locations can therefore be re-established in a short period of time and with marginal acquisition cost (as opposed to the situations such as the afore-mentioned study of bison). Such a rapid decay of past experiences is likely to be adaptive in dynamic landscapes akin to the one that roe deer experienced in this study (and which mimics dynamics in feeding site management by stakeholder outside of the experiment), as it allows animals to stay in tune with the spatio-temporal dynamics of their environment (Kraemer & Golding 1997), while reducing the physiological cost of memory storage and processing (Roth *et al.* 2010; Burns *et al.* 2011). An alternative explanation for the differing timescales of attribute memory between this study and Merkle et al. (2014) study of bison is that animals may use very recent information of a given patch profitability to determine its future quality (i.e., fast decay of the within-patch attribute memory; this study), but integrate information over longer temporal scales to assess the relative profitability of competing patch alternatives (i.e., slow decay of the between-patch attribute memory; (Merkle *et al.* 2014)).

More generally, our study demonstrates and quantifies the use of “where-what” memory by wild animals to guide foraging decisions in nature. Future research may further develop mechanistic models integrating episodic, or periodic (“where-what-when”) memory (Clayton & Dickinson 1998). For example, green-backed firecrown hummingbirds (*Sephanoides sephaniodes*) can achieve substantial energy gains by memorizing the spatial location (“where”) of high-quality feeders (“what”) and adjusting visit frequency (“when”) to nectar renewal dynamics (González-Gómez *et al.* 2011). In our experiment, a single perturbation of resource availability allowed to disentangle “where-what” memory from perception as main driver of movement, but multiple cycles, ideally of varying duration, would be needed to test episodic memory-based, mechanistic models.

Site familiarity is thought to provide fitness benefits in relation to foraging efficiency (Powell & Mitchell 2012) or predation avoidance (Gehr *et al.* in press), and to emerge from the re-visitation of known areas through spatial memory (Van Moorter *et al.* 2009). In large herbivores, observational studies have shown that animals select for previously-visited locations, such as bison (Merkle *et al.* 2014, 2017), caribou (Avgar *et al.* 2015), elk (*Cervus elaphus*) (Dalziel *et al.* 2008; Wolf *et al.* 2009), and feral hogs (*Sus scrofa*) (Oliveira-Santos *et al.* 2016). In such analyses, however, it remains difficult to ascertain whether animals revisit locations because of their intrinsic familiarity (i.e., familiarity effect) or because these locations are characterized by unknown, valuable environmental conditions (i.e., spurious familiarity effect *sensu* (Van Moorter *et al.* 2013)). Here, we found that roe deer strongly selected for their most familiar feeding site (M, by definition; see *Methods*) even after individuals had knowledge of equally profitable, alternate feeding sites (Fig. 2; Supplementary S1: Figure S1; Table 1). Because the identity of the familiar feeding site was not consistent across animals (i.e., a given site could be the most familiar and preferred by an individual, and avoided by another) it is unlikely that individual preference for a given feeding site resulted from location-specific, environmental conditions. While the adaptive value of familiarity has not been demonstrated here, the restoration of pre-disturbance patterns of resource use observed in this experiment cannot be explained by optimal foraging theory alone and supports the existence of site familiarity effects (Ranc *et al.* 2019).

Our analysis also revealed how environmental cues influenced roe deer foraging. Specifically, we found that roe deer visit to feeding sites, at short temporal scales, was strongly influenced by patterns of illumination: roe deer were most likely to attend feeding sites at dusk and during the night (Supplementary S1: Figure S1; Table 1), in accordance with the species circadian activity and movement patterns (Pagon *et al.* 2013). The high attendance at dusk compared to dawn is probably the result of the low food intake throughout the day, when feeding sites are avoided. The periodical patterns of feeding site attendance by roe deer resembles that of high-nutritional, but risky open landscapes such as agricultural fields (Bonnot *et al.* 2015; De Groeve *et al.* 2016, 2019), and is a clear example of recursion – a movement behavior tightly connected to spatial memory (Berger-Tal & Bar-David 2015). At longer temporal scales, roe deer significantly increased their use of feeding sites at low temperatures, consistent with the higher energetic demand of thermoregulation (Ossi *et al.* 2015, 2017). In the present study, the parameters associated with the environmental drivers of resource selection were highly conserved across the three cognitive hypotheses evaluated as evidenced by the high between-model overlap of the confidence intervals for illumination index variables and minimum daily temperature in Fig. 2. This implies that the effects of the considered environmental factors (i.e., temporal-dependency of resource selection) are relatively independent of the underlying cognitive mechanisms. Instead, long-distance (between-states) response to feeding site accessibility (absent in the perception-based model, by definition) and the preference for the manipulated feeding site (significantly different for omniscience-based model) varied considerably with the cognitive hypothesis considered.

Because foraging can be linked to fitness (Stephens & Krebs 1986), resource acquisition is considered to a primary driver of animal movement (Mueller & Fagan 2008). Although resource selection analysis has become a major tool in animal ecology (Boyce & McDonald 1999) that is used to inform conservation strategies (Chetkiewicz & Boyce 2009), the actual mechanisms through which wild animals interact with their surrounding landscape have not been elucidated. Further progress to connect animal behavioral decisions to individual performance (Gaillard *et al.* 2010), and ultimately, scale up to population dynamics (Morales *et al.* 2010) is contingent on our ability to uncover the mechanisms underlying animal movement in nature. Here, we have shown that memory-based movement models (specifically, a memory-based model of spatial transitions) parametrized using experimental data can successfully be used to quantify cognitive processes and to predict how animals respond to resource heterogeneity in space and time. The spatially-implicit modelling approach proposed in this study represents a stepping stone in the development of spatially-explicit, mechanistic models of animal movement and their parametrization using empirical data. By characterizing the spatial dimension of the interplay between memory and resource selection, such models would have the potential to shed light on the biological processes underlying home ranges in nature.

## Methods

### Study area

The study area, located in the north-eastern Italian Alps (*ca.* 50 km^2^; Autonomous Province of Trento), ranges between 600 and 1,000 m a.s.l and is dominated by mixed forest (> 80%). The climate is continental (mean daily temperature in January: 1.0 °C; in July: 21.0 °C; mean annual rainfall: 966 mm) with occasional snow cover. Roe deer is the most prevalent ungulate in the area (7-8 ind. km^−2^; ref. values from Autonomous Province of Trento Wildlife Office), and is selectively hunted between September and December. Supplemental feeding of roe deer is conducted year-round by private hunters at > 50 distinct feeding sites (FS; Supplementary S2: Fig. S1), typically shaped as hopper dispensers where corn can be accessed through a tray (Fig. 1, top row).

### Roe deer captures and collaring

Roe deer were captured using baited box traps near FS in winter (n = 15) and net drives in spring and fall (n = 3), and were fitted with GPS-GSM radio collars scheduled to acquire hourly GPS locations for a year, after which they were released via a drop-off mechanism. GPS acquisition success was extremely high (99.57 % during the experiment) and hence we did not interpolate missing fixes in the collected data. We collected data on 18 roe deer; 11 had collars for a single winter; two had collars that spanned two winters and five were recaptured and recollared for a second year, leading to a total of 25 animal-years (21 adults: 15 females, 6 males; 4 yearlings: 2 females, 2 males; see Supplementary S2 for details) in three consecutive years (n=4 in 2017, n=11 in 2018 and n=10 in 2019).

### Experimental design

Taking advantage of roe deer use of a focal, identifiable resource – the supplemental FS – we designed an *in situ* manipulation of resource accessibility for evaluating competing hypotheses pertaining to the processes governing roe deer foraging decisions. We created three successive experimental phases – pre-closure, closure and post-closure – by physically managing the accessibility of food at the FS. During the closure phase, we transitorily restricted the access of forage at manipulated FS (M) by placing wooden boards obstructing the tray (Fig. 1). Forage presence was maintained constant throughout the experiment at all FS. During the closure of FS, roe deer could sense corn through visual and tactile cues at short distances, as well as olfactory cues at both short and long distances.

During the *pre-closure phase*, we used roe deer movement data to identify M, defined as the most attended FS (hence, the most familiar) for each animal-year (see Ranc et al.(2019) for details). During the *closure phase*, we made corn at M inaccessible for a duration of about 15 days (min = 14.0 days, max = 18.1, mean = 15.5), depending on fieldwork constraints. We initiated the *post-closure phase* by restoring the accessibility of corn at M. During both pre- and post-closure phases, corn was available *ad libitum* to roe deer at M. All “alternate” managed FS (i.e., supplied at least once in the month prior to the experiment; A) had corn available *ad libitum* throughout the experiment.

The experiment was conducted in winter, when roe deer use of supplemental feeding is the most intense (Ossi *et al.* 2015, 2017). Animals were considered for the experiment after they revisited their capture location, used as indicator for the end of the post-capture response behavior. We ensured that co-occurring manipulations took place in separate areas to avoid potential interference. We defined animal-year (see above) as our sampling unit, on the assumption that the same individual may show an independent response to experimental manipulations in subsequent years as a consequence to varying internal (e.g., life-history), or external conditions.

### Spatial transition model

We developed a mechanistic spatial transition model to characterize the movements of roe deer between three distinct resource types, hereafter referenced to as states: the manipulated FS (M), alternate FS (A), and the matrix of natural resources, referred to as vegetation (V). We then characterized the movements of the roe deer between these states from their GPS locations. For M and A, we converted the FS point locations into areas by applying a buffer equal to the mean hourly step length of roe deer in our study area (i.e., 61.2 m). Previous analysis(Ranc *et al.* 2019) showed that the results of the analysis is not strongly affected by the choice of buffer size.

We derived the probability of moving to state **x**^*i*^(*t*) ∈ (*M*, *A*, *V*) at time *t*, for a given animal-year *i*, *p*^*i*^(**x**, *t*), from the attraction weight of state **x** in the previous hour, *w*^*i*^(**x**, *t* − 1):

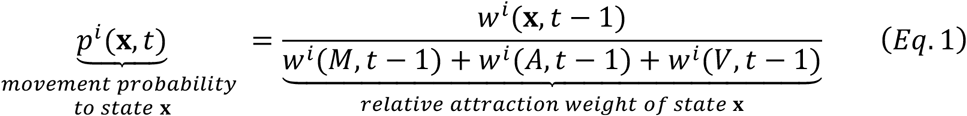

Because the movement probabilities were derived from relative attraction weights, we could simplify the model into an estimation of *w*^*i*^(*M*, *t* − 1) and *w*^*i*^(*A*, *t* − 1) by setting *w*^*i*^(*V*, *t* − 1) = 1.

Unlike other mechanistic models of resource selection (Van Moorter *et al.* 2009; Merkle *et al.* 2014), our model formulation does not only account for state-to-state movements (or patch-to-patch; e.g., transition from V to M) but is generalized to within-state movements as well (or residence; e.g., transition from V to V). This is achieved by defining the attraction weight of M and A as conditional on the state occupied by the animal at time *t*, **x**^*i*^(*t*).

### Within-state attraction

We defined *within-state attraction* as a function of the actual resource accessibility at M (*U*(*M*, *t*) = 1 in pre- and post-closure and *U*(*M*, *t*) = 0 during closure) or A (*U*(*A*, *t*) = 1 throughout the experiment), environmental covariates of FS use, *E*(*t*), and a population-level preference for M over A, *μ*:

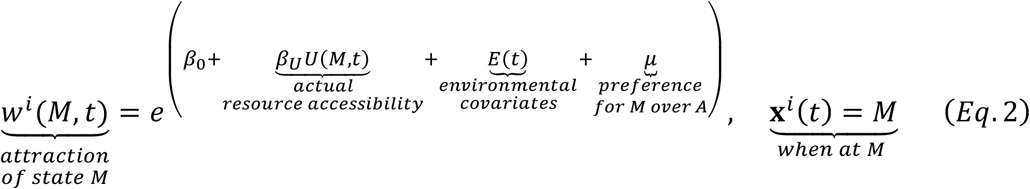

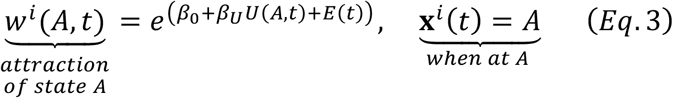

*Environmental covariates*, *E*(*t*) – In ungulates, and roe deer in particular, FS use is correlated with winter severity (Ossi *et al.* 2015, 2017), whose effect we approximated by using minimum daily temperature, *θ*(*t*). At shorter temporal scales, roe deer exhibit a strong circadian pattern in activity and movement behavior (Pagon *et al.* 2013), and in particular in their use of FS. For this purpose, we developed an illumination index derived from solar elevation, *I*(*t*), which approximates the sigmoidal shape of the log-transformed daily irradiance (Spitschan *et al.* 2017) (see Supplementary S3 for details). Because roe deer activity typically peaks during twilights and may differ between dawn and dusk, we included the rate of change of illumination, Ι_Δ_(*t*), and its absolute value, Ι_|Δ|_(*t*). The influence of environmental covariates on FS is given by:

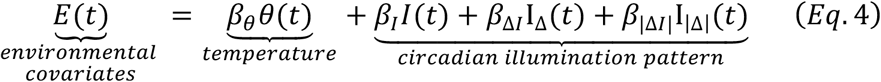

### Cognitive-dependence of between-state attraction

By contrast to within-patch attractions, the formulation of *between-patch attraction* in our model depends on which cognitive mechanisms roe deer use to evaluate the quality of distant resources (i.e., beyond current state). We formulated three competing cognitive hypotheses – omniscience, perception and memory – whose equations are detailed below. Because the equations characterizing the attraction of M and A only differ by the preference of M over A (*μ*, see Eq. 2) we present only the formulations for *w*^*i*^(*M*, *t*).

If roe deer possess a full knowledge of spatio-temporal resource dynamics i.e., *omniscience* (H1), between-state attraction depends on actual resource accessibility at M, *U*(*M*, *t*):

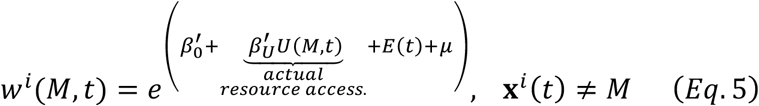

The above equation posits that the attraction of M for between-state attraction, independently of any covariates (i.e., the intercept), may differ from that of within-state attraction 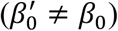. This conditionality on state occupancy, **x**^*i*^(*t*), allows to account for the high probability to remain within currently occupied state (i.e., 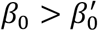), as indicated by the observed high serial correlation in the state time-series (see Ranc et al. (2019)). In addition, this formulation considers that roe deer response to changes in resource accessibility may affect movement (i.e., between-state transitions) and residence time (i.e., within-state transitions) differently such as 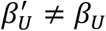.

If roe deer rely on *perception* alone (H2), they possess information (via olfaction) on resource presence at FS (constant at M throughout the experiment), but not on resource accessibility, which is manipulated (i.e., temporally-variable) at M. As a result, the between-state attraction equation for the perception model is not a function of *U*(*M*, *t*):

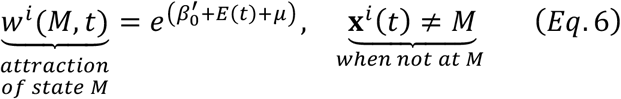

Alternatively, roe deer may rely on previous experience i.e., their memory, to guide foraging decisions (H3). Two different memory streams may be involved in decision-making: an *attribute memory* integrating previous experiences of resource quality to define the expected value of resource locations and a *spatial memory* encoding the spatial locations of resources (Fagan *et al.* 2013). Accordingly, roe deer movements should be influenced by expected resource accessibility – 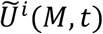 and 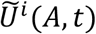 – instead of the actual resource accessibility – *U*(*M*, *t*) and *U*(*A*, *t*). We defined the expected resource accessibility as a temporally-weighted devaluation function of previous experience (Devenport & Devenport 1994). This formulation extends the exponentially weighted moving average of past experience (Killeen 1981; McNamara & Houston 1987), derived from the linear-operator model (Bush & Mosteller 1951), by accounting for the time interval between subsequent experiences and not only the serial order of experiences. We quantified the expected resource accessibility at M, 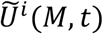, as the sum of experienced accessibility, *U*(*M*, *t*_*j*_), during all visits ***v***(*j* = 1…ϒ) at M that have occurred up to the current time *t*, and their associated times *t*_*j*_, weighted by their respective attribute memory, 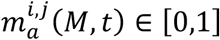:

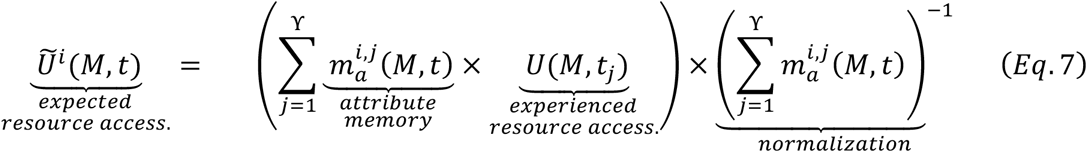

(see Merkle et al. (2014) for a similar formulation). The expected accessibility is updated at the end of each visit *j* such i.e., *t*_*j*_ satisfies **x**^*i*^(*t*_*j*_) = *M* and **x**^*i*^(*t*_*j*_ + 1) ≠ *M*. We modelled the attribute memory as an exponential decay function whose rate (0 ≤ *δ*_*a*_ ≤ 1) governs the devaluation of old experiences:

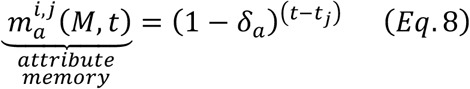

However, roe deer foraging decisions should not only rely on their capacity to integrate past experience of resource quality (attribute memory), but also on the ability to encode and retrieve spatial locations. To account for this process, we scaled FS attraction by a spatial memory weight, 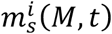:

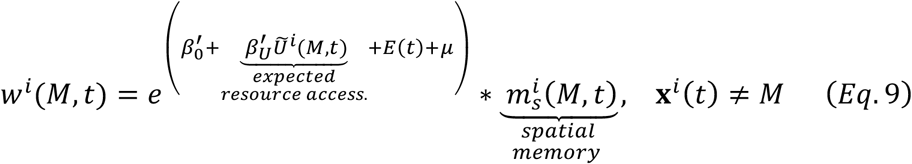

The spatial memory is maximum upon visit of the FS, and then decays exponentially with time since last visit (*t* − *t*_ϒ_), at rate *δ*_*s*_ (0 ≤ *δ*_*s*_ ≤ 1):

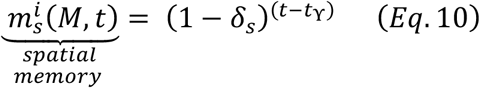

We chose to represent the temporal decay of attribute and spatial memory by negative (discrete) exponentials as this functional form of forgetting function is supported by substantial empirical evidence (Ziegler & Wehner 1997) and theoretical grounds (White 2001).

The spatial memory and expected resource accessibility values were initialized using the last encounter of M and A before the experiment onset (i.e., ***v***(*j* = 0)). For one individual, F4-2017, we did not have any recorded visit of A before the experiment and used her collaring date as visit event instead.

### Model parametrization and predictive ability

We estimated the model parameters through maximum-likelihood. The likelihood function for the parameter set 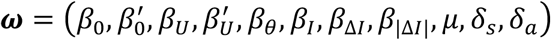 is given as:

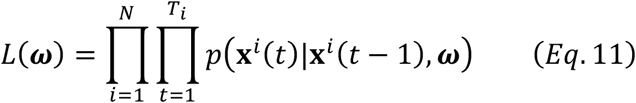

with *N* the number of animal-years (i.e., 25) and *T*_*i*_ the number of observations for animal-year *i* (missing GPS locations were omitted from the likelihood function). We used the particle swarm optimization algorithm (Poli *et al.* 2007) (PSO; see Supplementary S4 for details), a non-linear heuristic solver, to estimate the global minima of the log-likelihood function [*logL*(***ω***); i.e., the objective function]. We obtained 95% marginal confidence intervals (CIs) through an asymptotic normal approximation of the objective function in the neighborhood of the global minima. Because the objective function of the attribute memory decay rate was not well approximated by a quadratic function, we defined the CIs using a threshold of 1.92 log-likelihood value (as used for the ratio statistic) on either side of the global minima.

We evaluated the contribution of each variable by calculating the delta log-likelihood and delta Akaike information criterion (AIC) (Akaike 1973) of the reduced model (i.e., excluding the variable of interest) relative to the full model. For the memory model, we evaluated the contribution of spatial memory by setting δ_s_ = 0 (i.e., no decay), of attribute memory by setting δ_a_ = 1 (i.e., only the last experience is used to assess resource value), and of expected resource accessibility by setting 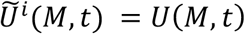 and 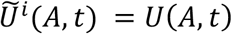 (i.e., expected equals actual resource accessibility, as in the omniscience model). We obtained the final models by dropping any covariates penalizing the AIC of the full models.

To evaluate the ability of the fitted mechanistic models to predict the movement behavior of roe deer during the experiment, we investigated whether they could capture the temporal dynamics in the rates of FS visit (V-to-M and V-to-A transitions), which summarize the general behavior of the system (Fig. 1). To this end, we compared the observed FS visit probabilities i.e., the transition probability matrix reporting p(M(t)|V(t-1)) and p(A(t)|V(t-1)), to associated predictions during the experiment. We obtained a temporal trend in transition probabilities from observed (discrete) transitions by calculating a running four-day mean.

Equations 1-11 were solved numerically in C++ and the parameters estimated using the PSO algorithm in MATLAB R2017b (MathWorks, Natick, Massachusetts, USA) using the Global Optimization Toolbox. The optimization ran on a computer cluster using the Distributed Computer Server (Sharma & Martin 2009). We calculated the illumination index (GeoLight package (Lisovski & Hahn 2012)), the CIs and produce effect size plots in R (R Development Core Team 2016).

## Acknowledgments

N. Ranc was supported by a Harvard University Graduate Fellowship and a Fondazione Edmund Mach International Doctoral Programme Fellowship. F. Cagnacci was supported by the Sarah and Daniel Hrdy Fellowship 2015-2016 at Harvard University OEB during part of the development of this manuscript. We are very grateful to all the people involved in the roe deer captures and fieldwork: M.B. Almeida, P. Bonanni, A. Corradini, J. De Groeve, K.W. Hansen, G. Lilli, S. Mumme, S. Nicoloso, D. Righetti, M. Rocca, M. Salvatori, T. Sforna, M. Sanchez and P. Semenzato. We thank J.W. Cain, P. Krastev (FAS Research Computing), A. La Fata, B. (Ölveczky, N. Pierce, and all the members of the Cagnacci lab and of the Moorcroft lab for their valuable suggestions. We also thank the Wildlife and Forestry Service of the Autonomous Province of Trento and the Trentino Hunting Association (ACT) for their support. Finally, we express our gratitude to the hunters of Albiano, Civezzano, Fornace and Trento Nord without whose help this study could not be conducted.

