## Supplementary Materials 1/2 for "Experimental evidence of memory-based foraging decisions in a large wild mammal"

**Supplementary S1: Circadian patterns in feeding site visit**

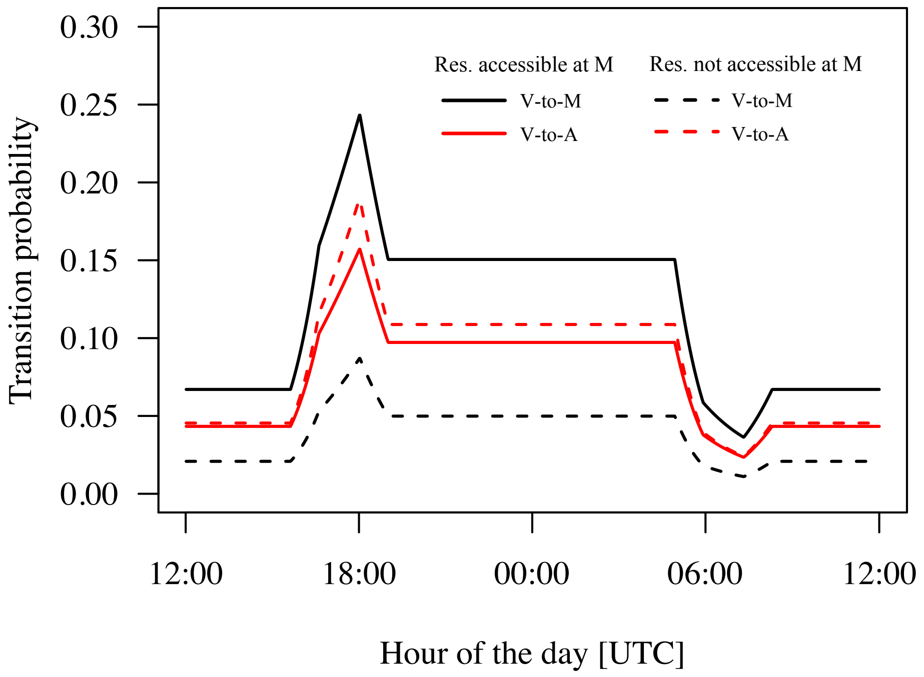


Figure S1. Circadian patterns in transition probabilities from vegetation (V) to either manipulated feeding site (M; V-to-M transition; black lines) or alternate feeding sites (A; V-to-A; red lines). The predictions are presented for resources accessible at M (during pre- and post-closure phases; solid lines); or inaccessible (during closure phase; dashed lines). The predictions were obtained from the reduced memory model with a maximum spatial memory value (i.e., full charge), no attribute memory (i.e., attraction based on last experience) and for the average minimum daily temperature recorded over the experiment (1.4°C). The conversion of illumination patterns to hour of the day is plotted for the mean date of the experiment (25$^{th}$ of February).

### Appendix S2: Animal captures and tracking

Between November 2016 and February 2019, we captured and marked 37 roe deer using wooden box traps baited with corn near FS in winter (n = 33) and net drives in spring and fall (n = 5). Of these captured individuals, 26 (yearlings and adults, or fawns captured after March) were fitted with GPS-GSM radio collars (VECTRONIC Aerospace GmbH; models GPS Plus, Vertex Plus or Vertex Lite). Captures and marking were performed complying with ethical and welfare rules, under authorization of the Wildlife Committee of the Autonomous Province of Trento (Resolution of the Provincial Government n. 602, under approval of the Wildlife Committee of 20/09/2011, and successive integration approved on the 23/04/2015). Nine individuals were recaptured in two separate years (n=7) or had data spanning two subsequent winters (n=2), thereby leading to a total of 35 animal-years (28 adults: 21 females, 7 males; 7 yearlings/fawns: 5 females, 2 male). Two collar batteries failed prior to this period. In addition, prerequisites for performing the experimental manipulation on an animal-year were: (i) spatial overlap between the animal-year movement trajectory and a FS, defined here as at least 10 relocations within a radius $l$ (mean hourly step length i.e., 61.2 m) of any managed FS, during a two-week period (i.e., the pre-closure) and (ii) possibility to alter the FS management, after explicit agreement with its private owner, which led to the exclusion of eight animal-years from the experiment. In light of the above considerations, **we retained 25 animal-years** (21 adults: 15 females, 6 males; 4 yearlings: 2 females, 2 males; n=4 in 2017, n=11 in 2018 and n=10 in 2019) for the experimental manipulation. Because one animal died (F4-2018) and another experienced a prolonged series of missing fixes (F28-2019) during the last phase of the experiment, we truncated two post-closure phases.

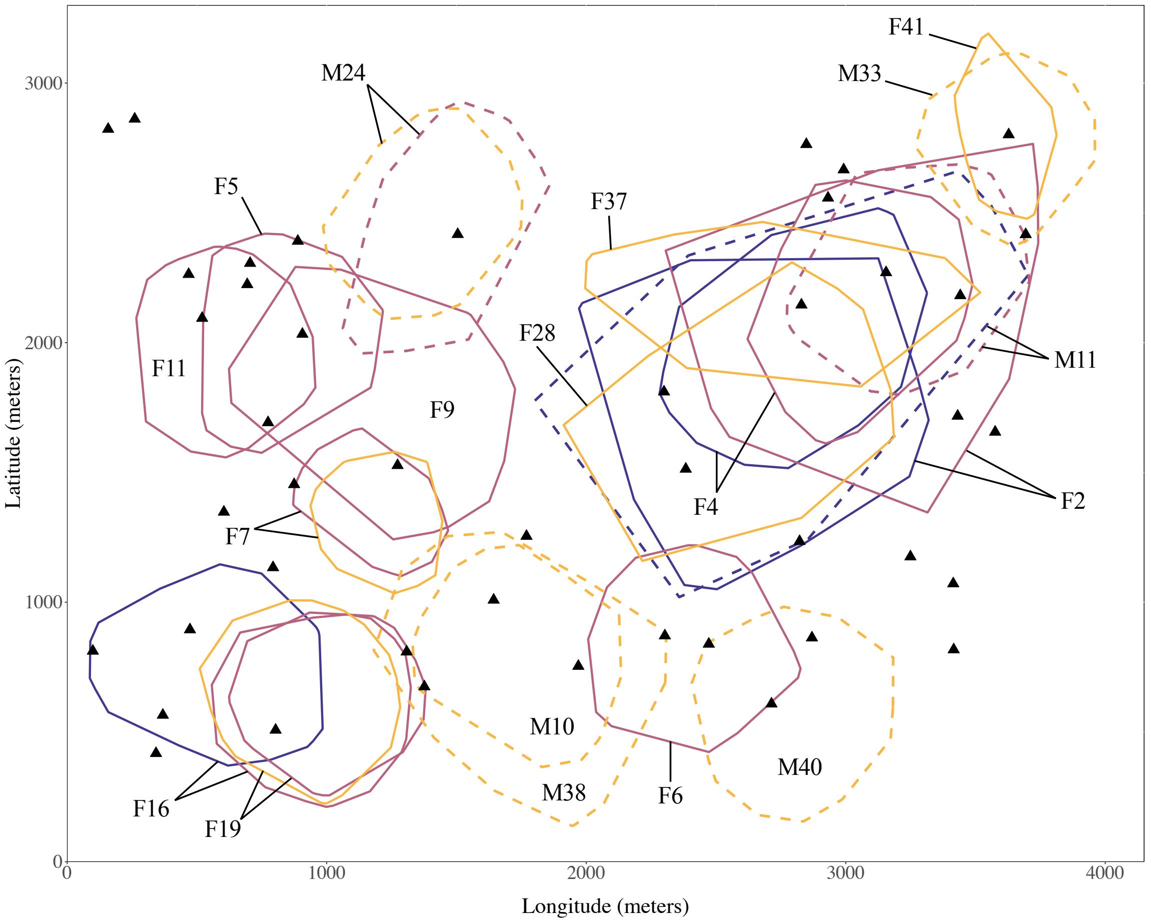


Figure S1. Spatial distribution of the roe deer included in the experiment. The area occupied by each twenty-five animal-years is plotted by a 95 % minimum convex polygon (2017: blue; 2018: burgundy; 2019: orange). Female home ranges are displayed as solid lines and males as dashed lines. All managed feeding sites (FS i.e., either M or A) are identified as black triangles.

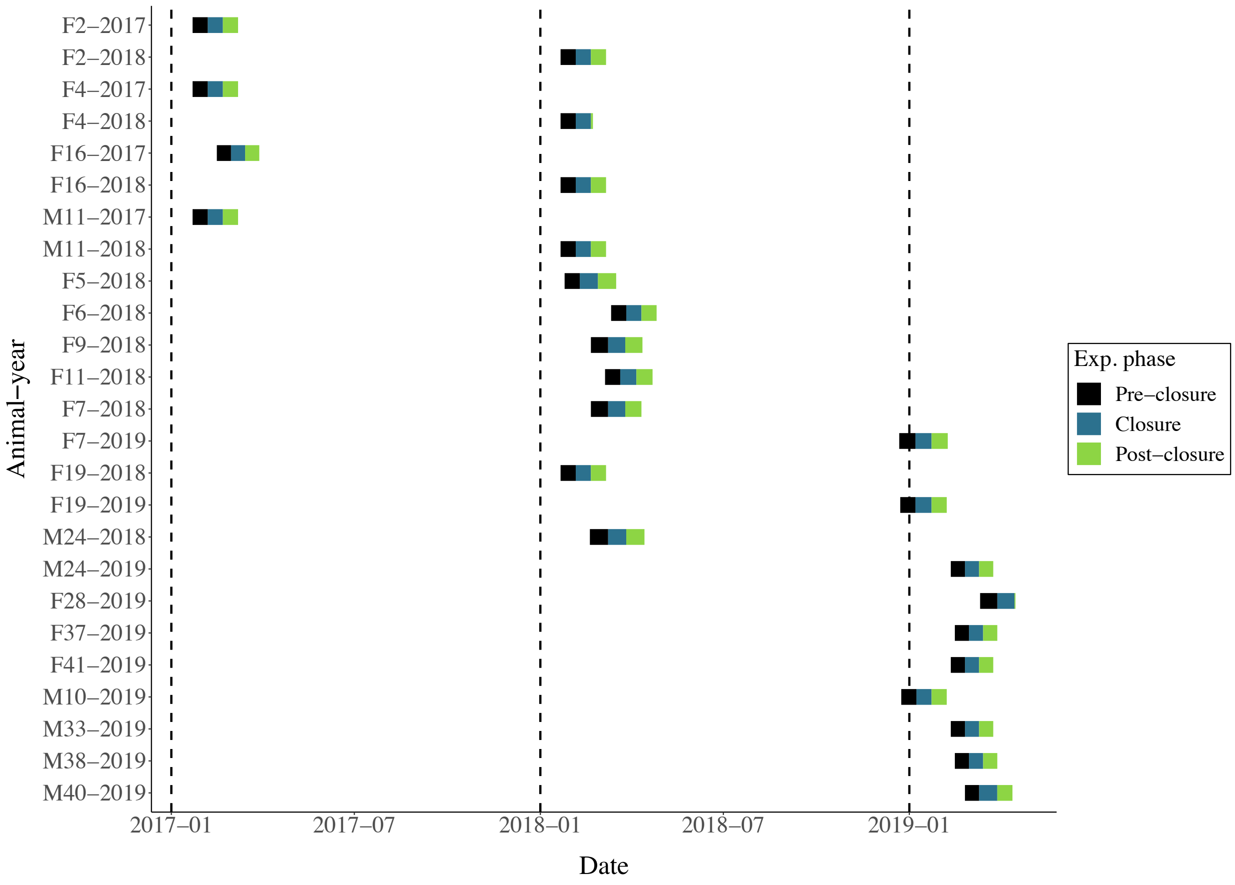


Figure S2. Monitoring history of the roe deer included in the experiment. To ensure comparability among animal-years, the initial excess positions for the pre-closure and closure phases and terminal excess positions for post-closure phase were truncated. The post-closure phases of F4-2018 and F28-2019 have been truncated due to a mortality case and a high proportion of missing fixes, respectively.

**Supplementary S3: Calculation of the illumination index**

Despite the overwhelming evidence that animal behaviour varies along circadian rhythms, few studies have explicitly accounted for the temporal-dependence of habitat selection. For example, roe deer activity patterns show very strong circadian signatures, being most active during twilights (Pagon *et al.* 2013) and selecting for cover during the day (De Groeve *et al.* 2016, 2019). Previous studies have used harmonics of time of day (Forester *et al.* 2009; Oliveira-Santos *et al.* 2016) to model complex interactions between circadian patterns and habitat selection. However, time of day may not be a suitable proxy for illumination (the underlying driver of circadian patterns) due to the confounding effect of seasonality. This issue becomes increasingly severe at latitudes away from the Equator, and for studies spanning several months. To address this shortcoming, other studies have developed a categorical time variable, typically night/day/twilight based on illumination patterns (Godvik *et al.* 2009; van Beest *et al.* 2012; Meisingset *et al.* 2013; De Groeve *et al.* 2016, 2019; Prokopenko *et al.* 2017).

In our analysis, we went further by modelling illumination as a continuous variable based on the solar zenith angle. The desired function should be minimum during the night and maximum during the day, when visible light is relatively constant, but should vary during the twilights. We defined the twilights as the temporal interval in which the sun is within 12° of the horizon. Our definition of twilight therefore includes the nautical twilight (the sun is between 12° and 6° below the horizon), civil twilight (between 6° and 0° below the horizon) and the beginning or end of day (between 12° and 0° above the horizon) for dawn and dusk, respectively. We set the value of our illumination index, $I(t)$, to 0 during the night, 1 during the day and a linear interpolation during the twilights (Fig. S1, top panel). The resulting function approximates the sigmoidal shape of the log-transformed daily irradiance obtained from empirical measurements (Spitschan *et al.* 2017). Because roe deer activity typically peaks during twilights and may be differ between dawn and dusk, we further derived the rate of change of illumination, $I_{\Delta}(t)$ (Fig. S1, central panel), and the absolute rate of change of illumination, $I_{|\Delta|}(t)$ (Fig. S1, bottom panel).

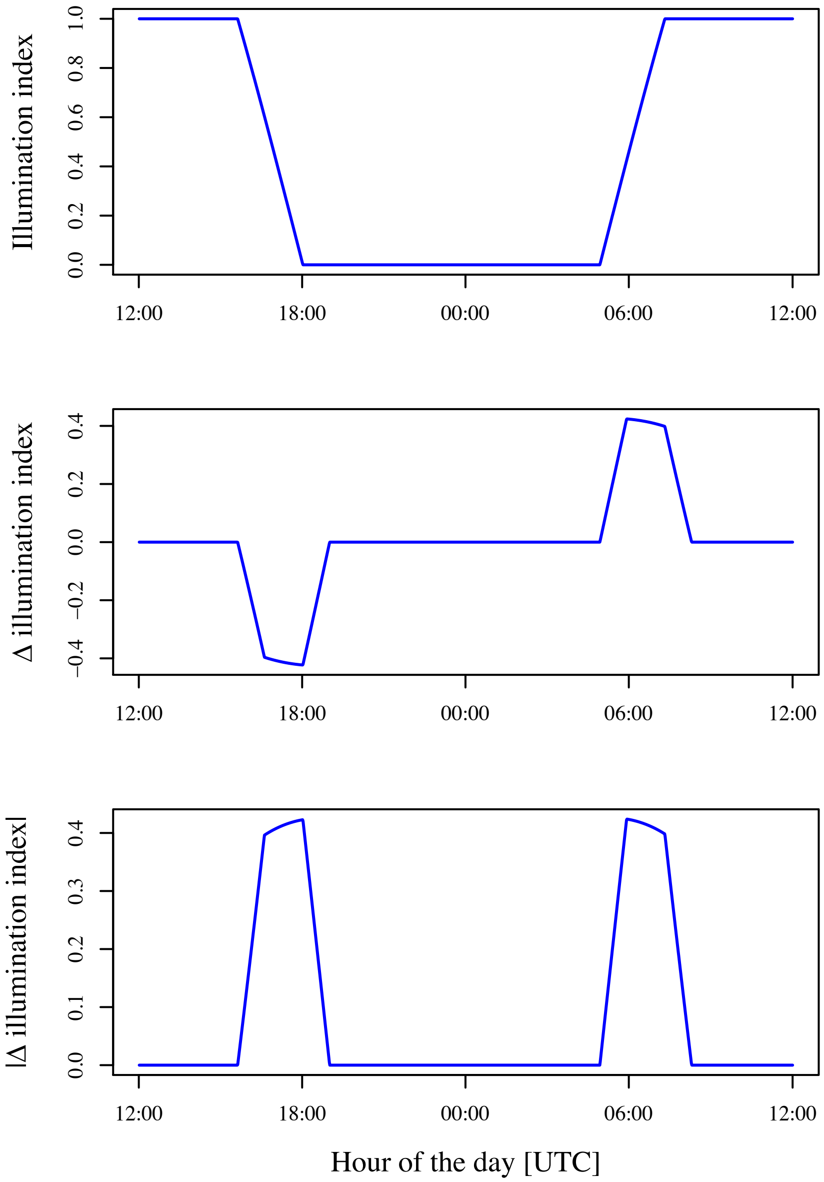


Figure S1. Illumination index ($I(t)$, top panel), its rate of change ($I_{\Delta}(t)$, central panel) and its absolute rate of change ($I_{|\Delta|}(t)$, bottom panel) as a function of hour of the day (UTC) for the mean date of the experiment (25$^{th}$ of February).
