## Supplementary Materials 2/2 for "Experimental evidence of memory-based foraging decisions in a large wild mammal"

### Appendix S4: Particle Swarm Optimization

Particle Swarm Optimization (PSO) is a non-linear heuristic algorithm using collective behavior to converge to a solution into the parameter search-space (see Poli et al.(2007) for a review). Given the relatively high number of dimensions of our problem, we initially chose to use a large swarm (180 particles) for a duration of 500 iterations, which revealed to be sufficient for our task. We did not define any a priori criteria of convergence and let the PSO reach the maximum number of iterations. We implemented a constriction PSO, which stabilizes the algorithm by dampening the velocity of the particles in the search space (Poli *et al.* 2007). We used the optimal settings described by Clerc & Kennedy (2002):

Constant, non-adaptive inertia ($\omega$) = 0.7298

Self-adjustment weight ($\emptyset_1$) = 1.49618

Social adjustment weight ($\emptyset_2$) = 1.49618

We used the algorithm available in the MATLAB Global Optimization Toolbox (MathWorks, Natick, Massachusetts, USA), with the default value for the Neighborhood fraction (0.25). To facilitate and increase the speed of the optimization convergence, we constrained the parameter search-space as follows:

$$\beta_0, \beta_0', \beta_U, \beta_U', \beta_\theta, \beta_I, \beta_{\Delta I}, \beta_{|\Delta I|}, \mu \in [-10,10]$$

$$\delta_s, \delta_a \in [e^{-20}, 1]$$
